# Screening of Potential Vaccine Candidate and Sera Based Diagnostic Markers from N-linked Surface Glycoproteins of *Entamoeba histolytica*

**DOI:** 10.64898/2026.07.24.740576

**Authors:** Santoshi Nayak, Mithu Baidya, Baisakhee Saha, Dipanwita Patra, Rashidul Haque, Sudip K. Ghosh

## Abstract

Pathogenesis inflicted by *Entamoeba histolytica* causes amoebic diarrhea and liver abscesses and is one of the leading causes of mortality from parasitic disease worldwide. Preventive therapeutics in the form of a vaccine could be highly effective at providing umbrella protection for a vulnerable community. In this study, a group of ten putative hypothetical surface N-linked glycoproteins was examined to assess their potential as vaccine candidates and/or diagnostic markers against amoebic intestinal colitis and liver abscesses. To evaluate the immunogenicity of putative surface glycoproteins, we used Entamoeba histolytica-infected patients’ sera from Bangladesh and then assessed the titer of antibody these glycoproteins elicited in patients by enzyme-linked immunosorbent assay (ELISA) and immunoblot. Based on this study, eight of ten surface glycoproteins were found immunogenic, as specific antibodies against these glycoproteins were detected in patients’ sera. Three of the immunogenic glycoproteins showed strong IgG antibody responses in both patients with intestinal amoebiasis and those with liver abscess. On the other hand, the other two glycoproteins showed the presence of specific serum antibodies exclusively in patients with amoebic liver abscesses, not in individuals with amoebic colitis. The remaining two glycoproteins showed more specific sera antibodies against ALA, but the preference was not very distinct. Immunolocalization with a specific antibody against the most immunogenic glycoproteins further confirmed their presence in the cell membrane. This differential immunogenicity of these glycoproteins in the two groups of patients qualifies them to become prospective sera-based diagnostic markers and also has the potential to become vaccine candidates for amoebiasis protection.

## INTRODUCTION

Intestinal colitis and liver abscess due to *Entamoeba histolytica* (Eh) claims about 100,000 deaths each year (1). Incidence of infection is more prevalent in tropical and sub-tropical countries in socio-economically disadvantaged communities having poor sanitary conditions (2, 3), of which children between ages of 2 to 5 are most vulnerable (4). Extensive studies in Bangladeshi children revealed that incidence rate of Eh related diarrhea is quite high at 0.08/child-year (5) but some children did obtain transient protective immunity after a natural Eh infection (6–8). This protective immunity against re-infection in children is correlated with secretory Gal lectin specific IgA antibodies in stool (7, 8). Since then serum specific anti-Gal/GalNAc lectin IgG is successfully used for clinical diagnosis of Eh infection (9) and is considered a very potential vaccine candidate (10, 11). In the urban slum of Fortaleza, Brazil where 25% of the people tested positive for anti-Gal/GalNAc lectin of *E. histolytica* and it was high at 40% for children aged 6–14 years (12). Similarly in a different study from South Africa it was shown that individuals having suffered from ALA with mucosal antibodies against Eh lectins showed higher protection against related *Entamoeba dispar*, a related species (13). Therefore, it seems humoral immune response do provide protection against both intestinal colitis and ALA and hence could be exploited to develop a potential vaccine. Other than Gal/GalNAc lectin various other protein were subsequently reported as a vaccine candidate among them is serine-rich *E. histolytica* protein (SREHP), whose characteristics like surface location, ability to induce adherence inhibitory antibody indicates it be a good vaccine candidate (14, 15). A protein worth mentioning will be a 29-kDa cysteine rich protein, which plays a role in inactivation of hydrogen peroxide, is found to be very immunogenic as antibodies recognizing this protein is said to occur in 80% of patient having amoebic liver abscess (ALA) (16). Other potential vaccine candidate reported are unique lipophosphoglycan on Eh surface (17, 18) cysteine proteinases like EhCP112 and EhADH112 (an adhesin) (19), and 30-kDa collagen binding protein (20) wherein these groups of molecule seem to provide fair resistance to ALA when studied experimentally in model organisms. Therefore, anti-amoebic antibody in serum could be a good indicator for infection and resilience of one’s immune system providing protection against recurring infection, but more importantly it can also adjudge immunogenicity of an antigen empirically. Apart from being a potential vaccine candidate few other surface lectins were also presented as possible diagnostic marker candidates. Of which EhChitinase and EhJacob2 are most notable as they were found to be polymorphic and could distinguish clinical isolate of Eh than other species (21, 22). Moreover, EhJacob2 was also found to be immunogenic and could successfully bind to anti-amebic patient sera (22).

Antigens well exposed on the surface of the pathogen help early recognition and protection by mucosal immune defenses and can prevent amoebiasis at its early onset. Therefore, most of the reported vaccine candidate till date is found to be membrane glycoproteins or lectins. Comprehensive cell surface N-glycoproteins identified by Carpentieri *et al*., (23) made known hundreds of surface N-glycoproteins, among them majority are annotated as hypothetical membrane proteins by Genome project analysis and are found to be abundant and are unique to Eh. These findings encouraged us to investigate if these putative surface glycoproteins can really stand out as promising vaccine candidates and diagnostic markers. Hence, in this present study we have heterologously expressed and purified few of these surface proteins and identified their reactivity with patient sera, who have suffered from either intestinal amoebiasis or ALA. Further by immunolocalization the putative antigens identified are found to be surface proteins.

## MATERIAL AND METHODS

### Animals

New Zealand white female rabbits which were used for polyclonal antibody production were purchased from an authorized dealer in Kolkata, West Bengal and were maintained at proper room temperatures. Institutional Ethical and Animal Care guidelines were adhered to during the experiments which are in agreement with “Committee for the purpose of control and supervision of experiments on animals (CPCSEA), Ministry of Environment and Forests, Government of India.

### Collection of patients’ sera

De-identified patient sera, received from five individuals with amebic liver abscess and five individuals with intestinal amoebiasis were used for the study were collected prior to the initiation of these studies from Bangladesh. The Ethical Review Committee of the International Centre for Diarrhoeal Disease Research, Bangladesh (ICDDR,B) reviewed and approved the design of the previous study under which these samples were obtained.

### Expression and purification of recombinant proteins and generation of polyclonal antibodies

Sequences for *E. histolytica* HM-1:IMSS surface N-linked glycoproteins are were retrieved form “AmoebaDB” (24). Since proteins under study are putative membranous hypothetical protein, therefore, we detected the presence of transmembrane helix by TMHMM (25). These regions of proteins were not considered for bacterial expression as they cause insolubility. GPI-anchor signals were identified with GPI-SOM (26). All the proteins were cloned in pET21a (+) vector, which were subsequently expressed in C43 (DE3) *E. coli* strain, suitable for membrane protein expression (27).Proteins were purified by Ni-NTA affinity chromatography (Qiagen, Germany) and the purity of proteins was analyzed in SDS-PAGE.

For primary immunization, purified proteins dissolved in PBS at a concentration of 2 mg/ml were emulsified with an equal volume of complete Freund’s adjuvant (CFA). Pre-immune sera were obtained before the immunization procedure. Rabbits were immunized subcutaneously at multiple sites along the spinal cord. Booster injections were administered with incomplete Freund’s adjuvant (IFA), at half the dose equivalent to that of primary. First booster was injected at 4^th^ week, followed by second booster on 6^th^ and the third booster on 8^th^ week. Blood was drawn from the marginal ear vein and the titers of antibodies were evaluated after every booster. Rabbit IgG antibodies were isolated from serum by ProteinA-sepharose (Invitrogen, USA).

### Immunoblot with patients sera

For assessing immunogenicity of the recombinant proteins by Western blotting, ∼3 µg each of purified recombinant proteins were separated in SDS-PAGE and transferred to a PVDF membrane. Membranes were probed with each patient’s sera (1:50 dilution) (Dhaka, Bangladesh) for overnight at 4°C. Membranes were then washed with for 15 min (three times) and then incubated with HRP-conjugated anti-human antibody (Sigma) (1:2000 dilution) for 1 hr followed by washing with PBS-Tween 20 like before. Immunoblot was finally detected using chemiluminescent kit (Luminata, Millipore, USA), as per manufacturer’s instruction. The presence of bound antibodies was scored as (+) and when absent it was marked as (−).

### Enzyme-linked immunosorbent assay for detection of specific anti-glycoproteins serum antibodies

Indirect ELISA was used to measure the specific anti-glycoproteins antibody titer in patient serum. Purified proteins were coated on to a 96-well F bottom microtiter ELISA plate (Tarson) in bicarbonate buffer such that each well contains approximately 10 μg of antigen. The plates were washed three times for 5 min with PBST (PBS containing 0.1% Tween 20 pH 7.4). To avoid nonspecific binding blocking was done using 3% bovine serum albumin in PBST and the plates were incubated for 1.5 hr at room temperature. The wells were again washed thrice using PBST and then incubated with 100 μl of patient sera with 1:1,000 dilutions in PBST having 1% BSA. After incubation for 2 hr at room temperature, each well was incubated with 100 μl of horseradish peroxidase-conjugated goat anti-human IgG (Sigma) (dilution 1:10,000) for 1 h at room temperature followed by thorough washing. Peroxidase activity on the ELISA plate was detected using 3, 3’, 5, 5’-tetramethylbenzidine (TMB)/H_2_O_2_ (Chromous Biotech, India) as enzyme substrate. Reaction was stopped with 1 M H_2_SO_4_ and theoptical densities were measured at 450 nm using Micro plate Reader (Multiskan, Thermo Scientific, USA). Values were corrected by subtracting negative control value, which was measured from the serum sample collected from a healthy individual donor with no amoebiasis history.

### Immunolocalization of the antigenic glycoproteins

*E. histolytica* **c**ells were harvested and then washed thrice in 1X PBS and then fixed with 4% paraformaldehyde for 10 min in ice. The fixed cells were then permeabilized with 0.01% Triton X-100 for few seconds. Cells were then stained with purified specific anti-glycoprotein antibody raised in rabbit for 1 hour at room temperature having 1:100 dilution followed by anti-rabbit IgG TRITC conjugated antibody (1:500). Stained cells were visualized by confocal microscopy (FV 1000, Olympus, Japan).

## RESULTS

### N-linked glycoproteins under study are predicted membrane bound surface proteins

Over more than two hundred surface glycoproteins have been reported by Carpentieri et al., (23) out of which ten hypothetical proteins are been chosen (preferable with one transmembrane domain) for the current study as the biological relevance of these vast pool of proteins are still unknown. All the hypothetical proteins were named as Eh surface protein followed by a number according to ascending order of their accession number in Amoeba DB (Fig 1A). To ascertain if these proteins are surface bound, they were checked for predicted transmembrane domains and glycophosphatidylinositol (GPI) anchor site, which will hook this proteins to the membrane. Analysis of these proteins on TMHMM server (25) reveals that all the Eh surface N-linked glycoproteins (EhSGP) have transmembrane helices at the C-terminal domains. However, only EhSGP7 has got transmembrane helices at both C and N-termini. Out of all the proteins EhSGP4, 6 and 9 have major portion of the protein tethered inside to the cell cytosol whereas rest of the proteins are found to be anchored to the membrane from outside of the cell membrane. On GPI prediction with GPI-SOM (26), only one glycoproteins (EhSGP10) was found to have GPI anchorage site both at the C-terminal end and are predicted to be hooked to the cell membrane from the cell exterior (Fig. 1a). To check their immunogenicity all the glycoproteins are heterologously overexpressed in bacteria bereft of their transmembrane helix domain and rare codon stretch (Fig. 1B).

**FIG 1:**
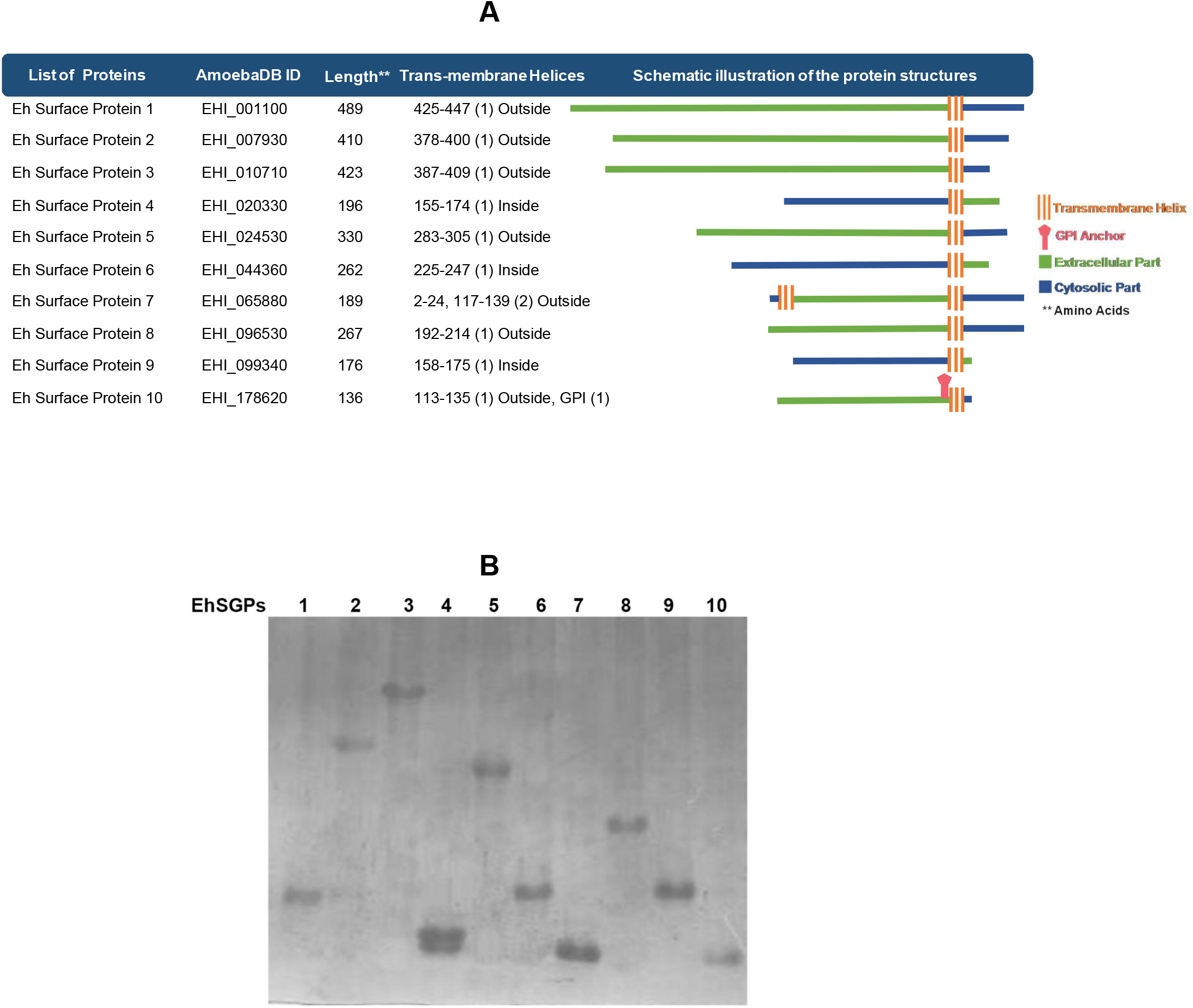
Expression and purification of recombinant glyoprotein fragments. (A) Schematic representation of *Entamoeba histolytica* surface glycoproteins under study, indicating transmembrane-helix, GPI anchors, and membrane positioning of the glycoroteins. (B) Coomassie-stained SDS-PAGE showing purified EhSGPs

### Eh surface glycoproteins show varied immunogenicity to intestinal colitis and ALA

To screen and assess immunogenicity of these surface glycoproteins specific serum IgG against individual protein were detected using immunoblot and ELISA. Immunoblot identified that anti-EhSGP1and 6 serum IgG were present in four out of five patients whereas serum IgG against EhSGP 9 were present in all the five patients having intestinal amoebiasis. Very poor response was observed for EhSGP 3 and 4 in intestinal amoebiasis patients’ sera with only one out of five individuals having sera IgG against these glycoproteins. Sera isolated from ALA patients showed presence of anti-EhSGP3 and 7 in all five individuals while anti-EhSGP4, 5 and 9 were found in four samples and 3 patients were detected for EhSGP6 IgG (Table 1). For relative quantification of sera anti-EhSGPs titer in each group of patients, ELISA test was carried out taking serum IgG from an individual having no amoebiasis history as blank. On quantification it was estimated that the highest amount of titer were found against EhSGP9, followed by 6, 1, 4 and 3 in group of patients suffering from intestinal amoebiasis while for the rest of the glycoproteins IgG titer was insignificant compared to normal individuals (Fig. 2A). In other group of patients with ALA, highest titer of specific EhSGP-IgG antibody were seen against EhSGP7, followed by 5, 3, 9, 4, 1 and 6 while for the rest the titer values were far less pronounced (Fig. 2B). These results ascertain that few of these surface glycoproteins have quite elevated serum reactivity and hence could be good candidates as vaccine and/or diagnostic marker.

**TABLE 1:**
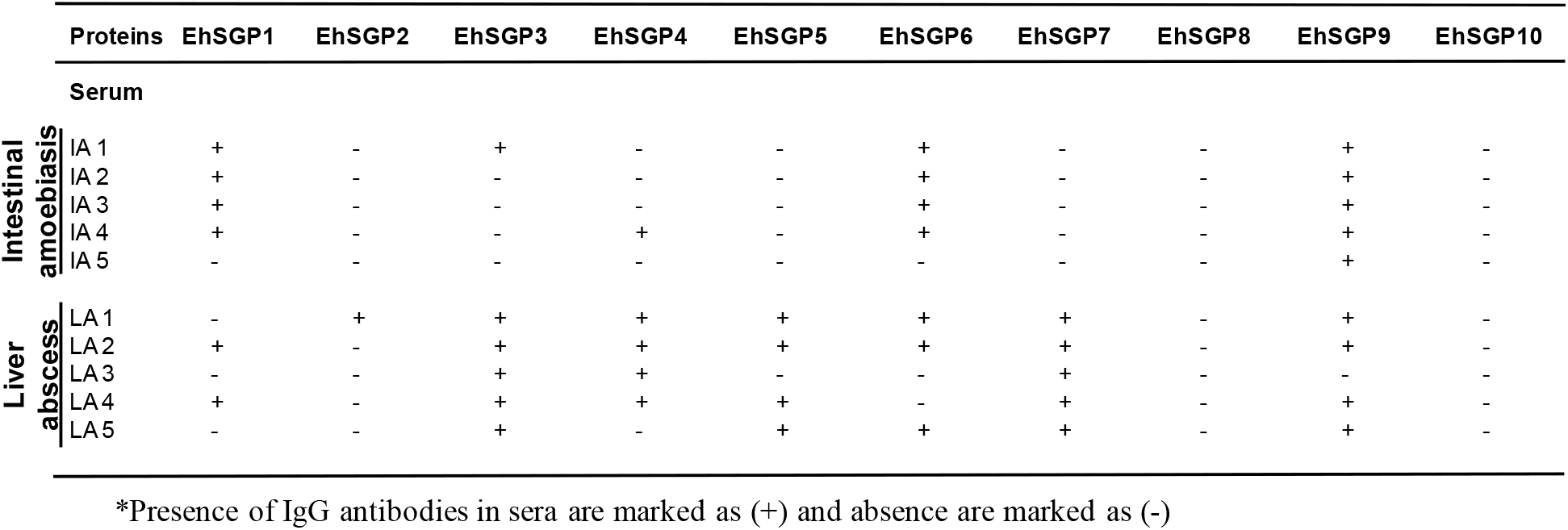
Prevalence of anti-EhSGP IgG antibodies in patients sera of Bangladesh as determined by immunoblot.

**FIG 2:**
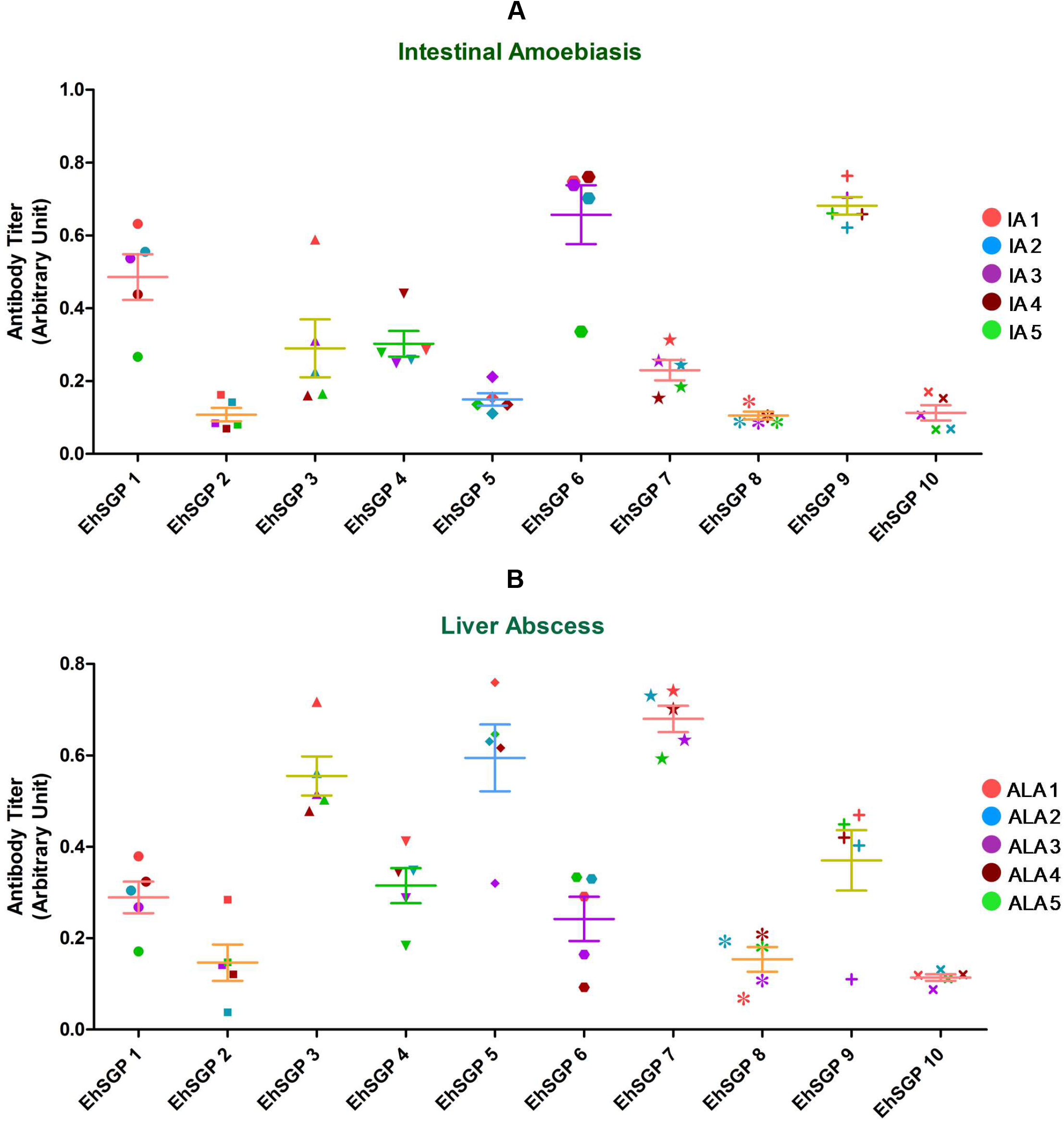
Estimation of srea anti-EhSGP-IgG antibodies titer by ELISA. Results demonstrating estimated presence of specific anti-EhSGP-IgG antibodies in sera of (A) intestinal amoebiasis patients and (B) amoebic liver abscess patients. The titer value is estimated in triplicate for each serum sample for individual recombinant EhSGPs and the mean values were normalized with healthy individual’s sera with no amoebiasis history. IA represents intestinal amoebiasis patients’ sera and ALA stands for sera from patients having amoebic liver abscess.

### Antigenic glycoproteins are localized on to the membrane of *Entamoeba histolytica*

Based on the concentration of sera antibody seven glycoproteins (EhSGP1, 3, 4, 5, 6, 7, 9) were found to be potential vaccine candidate and therefore, specific antibodies were generated against each of them in rabbit for assessing their immunolocalization. On microscopic analysis, EhSGP 1, 3, and 7 were found to be surface stained without cell permeabilization, which infers that the proteins were localized on the cell exterior. On the other hand, EhSGP 4, 5, 6 and 9 stained only on cell permeabilization indicating their membrane localization towards the cytosolic side. Although localized on the membrane, EhSGP 4 and 9 was also seen as distinct puncta in the cytosol. However, largely all the antigenic N-linked glycoproteins were found to be on the membrane of Eh as predicted.

## DISCUSSION

Till date, the only effective drug for amoebiasis treatment is metronidazole (28, 29). Very often these drugs are overused and hence clinical resistance to metronidazole could be disastrous as some laboratory strains are shown to have grown resistance against metronidazole on systemic overexposure (30). Moreover, improved sanitation and hygiene to exist in developing countries in foreseeable future is an unrealistic scenario. Therefore preventive vaccine can ensure check on its spread. Fortunately, *Entamoeba* has fairly simple life cycle compared to other protozoan parasite and has only one definitive host i.e. human without any intermediate vector. Moreover, it also remains extracellular inside the host which makes it relatively feasible to develop effective vaccine for providing protection to a population *en masse*. Finally, though there are limited evidences (7, 8) it has been found that prior infection imparts some immunity, which is very encouraging.

Attenuated *E. histolytica* were used for vaccination and was found successful in some of the animal studies (31). However, using killed or inactivated whole organisms as vaccine has always remained controversial. Therefore, focus has shifted to use of recombinant *E. histolytica* antigens. Though a plethora of surface glycoproteins were reported as vaccine candidates, but extensive surface glycoprotein’s immunogenicity profiling was never attempted. Here we have taken ten surface N-linked glycoproteins from *E. histolytica* to assess their immunogenicity in sera of patients having intestinal colitis and ALA. Out of ten glycoproteins under study, only 7 showed specific sera IgG titer against the respective antigen significantly. Interesting findings that underlined this study was that the proteins showed differential immunogenicity to intestinal amoebiasis and ALA sera. Three of the immunogenic glycoproteins (EhSGP1, 6 and 9) showed presence of specific antibodies in patients having both intestinal amoebiasis and liver abscesses. On the other hand, specific sera antibodies against other two glycoproteins (EhSGP 5 and 7) were exclusively found in patients having amoebic liver abscesses but not in individuals having amoebic colitis. The rest of the two glycoproteins (EhSGP 3 and 4) showed more specific sera antibodies in ALA patients while mild presence was seen in intestinal amoebiasis patients. However, other three (EhSGP 2, 8, 10) recombinant proteins showed least immunogenicity in any patients sera (Table 2). Another noteworthy observation was that cytosolic faced membrane proteins, EhSGP 5, 6 and 9 (Fig. 3), were found to be more immunogenic in patients having intestinal amoebiasis (Fig. 2A). The same recombinant proteins were also shown to have some reactivity in ALA sera. On the other hand, EhSGP7, which is hooked into the membrane from outside (Fig. 3) was present specifically in ALA patients only (Fig. 2B), but not in patients having intestinal colitis. Similarly, other surface proteins which are tethered to membrane from outside like surface glycoproteins EhMSP-1 (32), the serine-rich *E. histolytica* protein (SREHP) (15, 33), 29-kDa cysteine-rich thiol-dependent peroxidise(16), a lipoproteophosphoglycan (17, 18) and D-galactose/N-acetyl-D-galactosamine-specific amebic adherence lectin (Gal lectin) (33–35) were also found to be immunogenic in ALA patients as well as rendered protection against ALA in rodent models. So, further animal model based study is imperative to ascertain the protective ability of newly found antigenic glycoproteins against ALA so as to qualify them as vaccine candidate. Since EhSGP 5 and 7 are differentially immunogenic to intestinal amoebiasis and ALA, therefore, they have potential use in clinical serodiagnosis of ALA. Also, EhSGP 6 and 9 though not specific to liver abscess can also have the potential to be used as diagnostic marker for amoebic infection like Gal/GalNAc lectins (9, 36). Though limited numbers of patient’s sera were used for this pilot project, further studies using large sample size from different country can establish this early trend, which subsequently can qualify both as diagnostic marker and as protective vaccine. Though many potential vaccine candidates are under extensive study, yet no single clinical vaccine for human consumption is available, which thereby creates space for new entrants to contest the race.

**TABLE 2:**
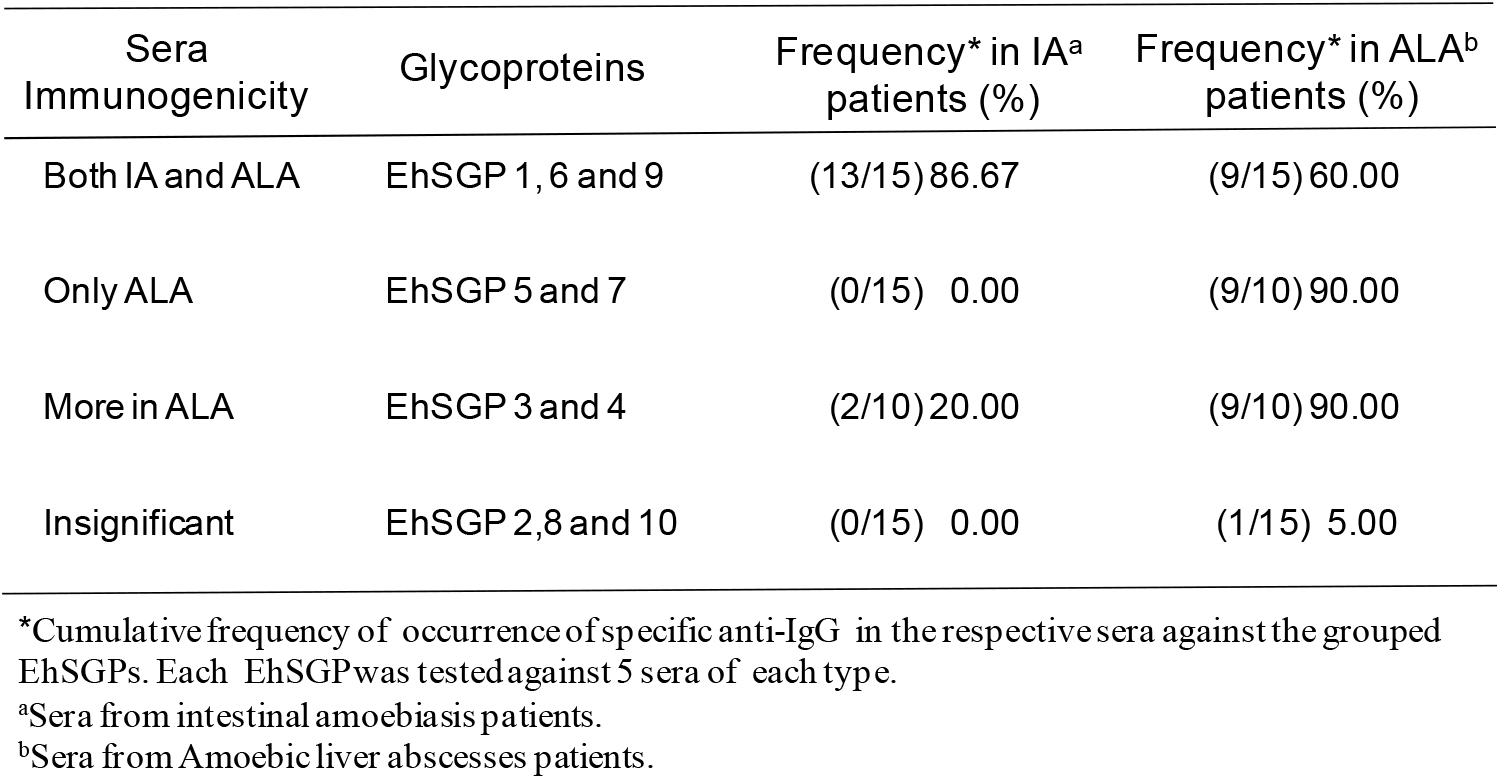
Sera specificity of EhSGPs as determined by immunoblotting.

**FIG 3:**
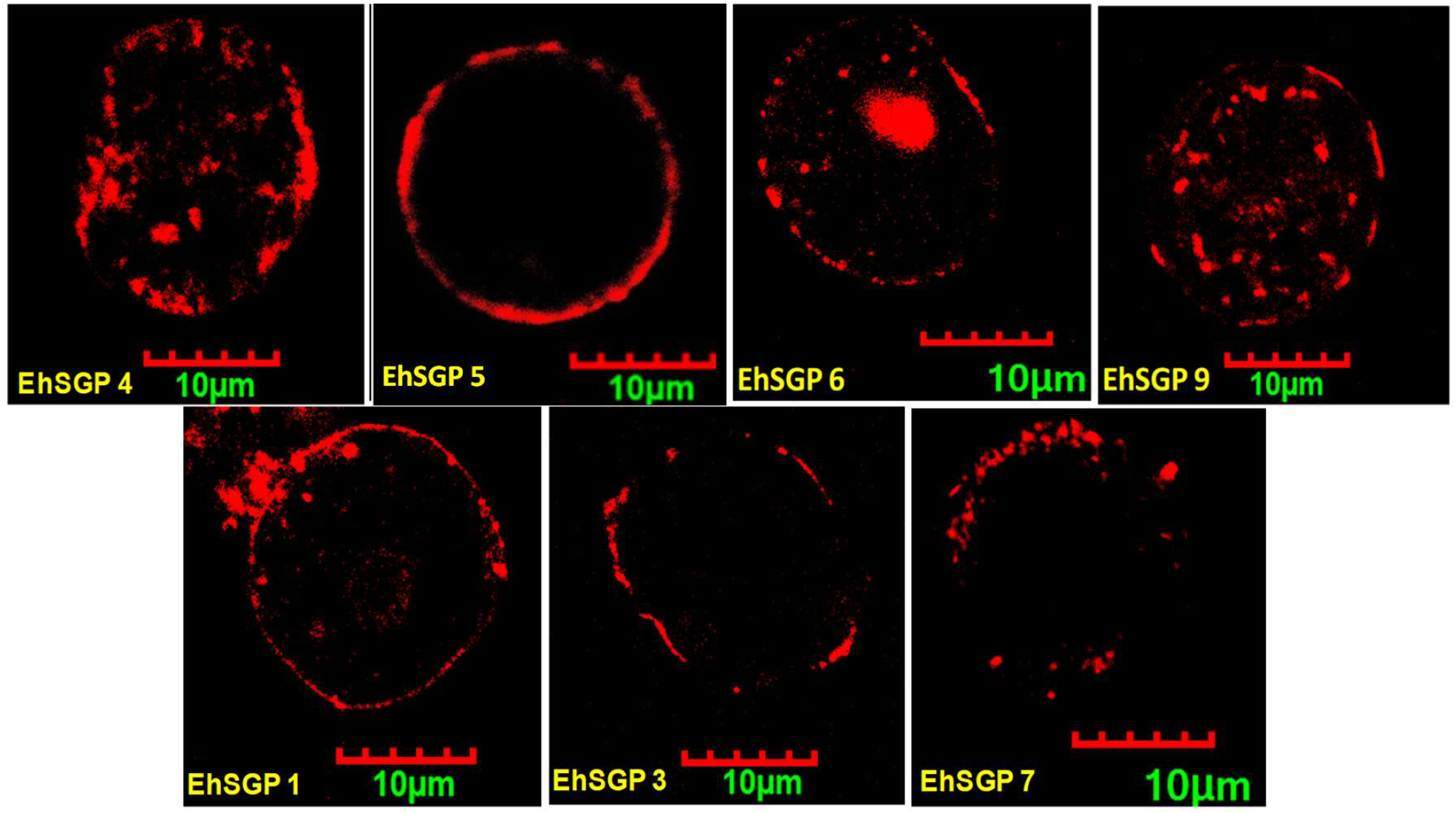
Localization of EhSGPs. Specific antibody generated against the very immunogenic recombinant EhSGPs in rabbit were used for immunolocalization. On confocal microscopy it was found that EhSGP 4, 5, 6 and 9 were stained only when cells were permeabilized with Triton X-100 and therefore, it is inferred that they are localized towards cytosolic side of the membrane. On the other hand, specific antibodies against EhSGP1, 3 and 7 could easily stain the respective EhSGPs without permeabilization indicating the proteins are localized on the surface of the membrane outside the cell.

## ACKNOWLEDGEMENTS

We are most grateful to the DST-FIST for the confocal facility at the Indian Institute of Technology, Kharagpur, India. This work was supported by MHRD, and Ministry of Health and Family Welfare, Govt. of India. Grant Number F. No.: 3-18/2015-TS-TS.I, Dt. 29-11-2016. S.N. is grateful to CSIR and M.B. is thankful to UGC for the award of a personal fellowship DP is thankful to IIT Kharagpur and BS to MHRD for Post fellowship.

## DECLARATION

All the authors declare there is no conflict of interest.

